# Ancient Fish Lineages Illuminate Toll-Like Receptor Diversification in Early Vertebrate Evolution

**DOI:** 10.1101/2023.04.05.535752

**Authors:** Kara B. Carlson, Cameron Nguyen, Dustin J. Wcisel, Jeffrey A. Yoder, Alex Dornburg

**Author notes:** Corresponding author at: Alex Dornburg, Department of Bioinformatics and Genomics, University of North Carolina at Charlotte, Charlotte, North Carolina, USA.

## Abstract

Since its initial discovery over 50 years ago, understanding the evolution of the vertebrate adaptive immune response has been a major area of research focus for comparative geneticists. However, how the evolutionary novelty of an adaptive immune response impacted the diversity of receptors associated with the innate immune response has received considerably less attention until recently. Here we investigate the diversification of vertebrate Toll-like receptors (TLRs), one of the most ancient and well conserved innate immune receptor families found across the Tree of Life, integrating genomic data that represent all major vertebrate lineages with new transcriptomic data from Polypteriformes, the earliest diverging ray-finned fish lineage. Our analyses reveal TLR sequences that reflect the 6 major TLR subfamilies, TLR1, TLR3, TLR4, TLR5, TLR7, and TLR11, and also currently unnamed, yet phylogenetically distinct TLR clades. We additionally recover evidence for a pulse of gene gain coincident with the rise of the adaptive immune response in jawed vertebrates, followed by a period of rapid gene loss during the Cretaceous. These gene losses are primarily concentrated in marine teleost fish and synchronous with the mid Cretaceous anoxic event, a period of rapid extinction for marine species. Finally, we reveal a mismatch between phylogenetic placement and gene nomenclature for up to 50% of TLRs found in clades such as ray-finned fishes, cyclostomes, amphibians, and elasmobranchs. Collectively these results provide an unparalleled perspective of TLR diversity, and offer a ready framework for testing gene annotations in non-model species.

## Introduction

Over the past decade, the accelerating pace of genome-sequencing has given rise to unprecedented insights regarding the macroevolutionary trends that orchestrate the evolution of gene repertoires across the Tree of Life (Hansson and Stensmyr 2011; McGowen et al. 2014; Musilova et al. 2019; Dornburg et al. 2021a). While cataloging the interspecies diversity of gene repertoires has the potential to reveal the role of gene gains and losses in promoting the evolution of functional novelty, large-scale comparative investigations into the diversification history of immune receptors in vertebrates have only recently begun to appear (Hughes 2010; Montaño et al. 2011; Langevin et al. 2013; Liu et al. 2020). The molecular basis of immunity in vertebrates is unusual and often divided into two dominant motifs: adaptive and innate immunity. Like other metazoans, every vertebrate possesses a germ line encoded innate immune response that forms a front line of defense against pathogens (Buchmann 2014; Scholz et al. 2022). In addition, jawed vertebrates (Gnathostomata) have evolved an adaptive immune response that allows for the creation of somatically recombined, pathogen specific responses which are retained to protect against future infection (Litman et al. 2010; Kaufman 2018). The early evolutionary history of the vertebrate adaptive immune response has been the subject of numerous investigations (Cooper and Alder 2006; Litman et al. 2010; Boehm and Swann 2014; Morales Poole et al. 2017; Kaufman 2018). Unfortunately, the evolutionary dynamics of innate immune receptors following the origin of the vertebrate adaptive immune system remains poorly understood.

The vertebrate adaptive immune response is heavily dependent on the proteins encoded by the major histocompatibility complex (MHC) and interacting receptors that distinguish self from nonself (Litman et al. 2010; Wilson 2017; Kaufman 2018). Since the early studies of antibody function (Edelman and Gall 1969), numerous immunological investigations have defined the association between adaptive immune receptors [e.g., antibodies and T-Cell Receptors (TCRs)] and transmissible as well as autoimmune disease outcomes (Cooper and Alder 2006; Murphy and Weaver 2016; Flajnik 2018). While the evolution of this mode of immunity has had a tremendous effect on the coevolutionary arms race between vertebrate hosts and their pathogens (Nourmohammad et al. 2016; Laanto et al. 2017), the diversity of vertebrate innate immune receptors has historically received considerably less attention (Cunliffe 2006). A flurry of studies have cast new light on this neglected topic, revealing an immense scope of sequence and gene family diversity we are only beginning to understand (Langevin et al. 2013; Dornburg et al. 2021b; Dornburg and Yoder 2022), as well as surprising key interactions between innate and adaptive immune responses (Clemente et al. 2016; Gianchecchi et al. 2018). Further, the realization that innate immune responses can be trained has led to a call for fundamental change in our perspective of immunological memory (Netea et al. 2019). It is becoming clear that rather than corresponding to two distinct systems of defense, adaptive and innate immune responses are part of an integrated evolutionary continuum of host defenses (Boehm 2012; Netea et al. 2019). As such, studies of innate immune receptor families that span all vertebrates are essential for providing insights into the mode and tempo of early vertebrate immune evolution. Moreover, such studies are vital for providing the historical context necessary to define homology between receptors and translate fundamental findings between model organisms and humans.

The Toll-like receptor (TLR) family is an ideal model for investigating the evolutionary dynamics of vertebrate innate immune receptors. TLRs are well-studied pattern recognition receptors (PRRs) and one of the most ancient and well conserved innate immune receptor families found across the Tree of Life (Leulier and Lemaitre; Gallucci and Maffei 2017). In invertebrates, TLRs serve multiple developmental and immune specific roles and can be present in the hundreds. In contrast, vertebrates possess only a fraction of this TLR phylogenetic diversity, with their known functions restricted mainly to the recognition of conserved pathogen-associated molecular patterns (PAMPs) and certain self-derived molecules from damaged cells, referred to as damage/danger-associated molecular patterns (DAMPs). TLR-mediated recognition of PAMPs plays crucial roles in the defense against fungal, bacterial, viral (including SARS-CoV-2 (Loske et al. 2022)), and parasitic infection, while recognition of DAMPs aids in the recognition of damaged or diseased cells (Roach et al. 2005; Kawasaki and Kawai 2014; Wcisel et al. 2017; El-Zayat et al. 2019; Liu et al. 2020). Analyses of the growing number of reference genomes indicate that TLR phylogenetic diversity is lower and functionally more restricted in vertebrates relative to multiple hemichordate, echinoderm, and select invertebrate lineages (Buckley and Rast 2015; Nie et al. 2018; Liu et al. 2020). However, sequencing of genomes from jawless vertebrates (agnathans) has revealed that major changes in TLR gene diversity predate the rise of MHC based adaptive immunity in jawed vertebrates. Compared to agnathans, numerous gnathostome lineages also possess an expanded repertoire of TLRs that varies dramatically between lineages (Liu et al. 2020; Carlson et al. 2022). For example, a number of model teleost fish species possess twice the number of TLRs as mammals, and until recently it was assumed that this disparity in TLR diversity was a consequence of paralog retention following the teleost genome duplication events (Sundaram et al. 2012; Solbakken et al. 2016; Khan et al. 2019; Smith et al. 2019). The sequencing of genomes from the sister lineage to teleosts, Holostei (spotted gar and bowfin), overturned this hypothesis revealing equivalent TLR diversity in lineages without a duplicated genome (Wcisel et al. 2017; Thompson et al. 2021). This example from teleosts illustrates the importance of placing the evolutionary history of TLRs into a deeper phylogenetic context and also raises a general question. What molecular mechanisms gave rise to the asymmetric patterns of TLR gene diversity maintained across vertebrates?

Several hypotheses could explain the evolution of vertebrate TLRs. On the one hand, the rise of adaptive immunity may have led to a further contraction of TLR diversity with observed levels of TLR diversity reflecting apical lineage-specific losses or gains across vertebrate clades. Such a pattern would be consistent with general evolutionary trends observed across metazoan gene families (Fernández and Gabaldón 2020), and also correspond with hypotheses that invoke pathogen driven diversification of TLRs (Loker 2012; Khan et al. 2019). On the other hand, it is possible that an early burst of TLR diversity early in the evolution of jawed vertebrates was catalyzed by the origin of the adaptive immune system. Under this hypothesis, the bulk of vertebrate TLR diversity would have arisen within a few early vertebrate lineages, reflecting neofunctionalization of early vertebrate TLR paralogs in concert with the diversification of new interacting partners managing pathogenic pressures. A recent investigation of TLRs across Metazoa suggested that several vertebrate TLRs do indeed have deep evolutionary origins (Liu et al. 2020), though lack of genomic resources from key early diverging actinopterygians precluded a definitive assessment of TLR origins for half of all living vertebrates. Given this conflicting evidence, the extent to which these related hypotheses explain TLR diversification remains unclear.

Here we generate novel transcriptome sequence data from Senegal bichir (*Polypterus senegalus*), a species belonging to the earliest diverging clade of actinopterygians (Polypteriformes) (Near et al. 2014). We integrate this data with newly available vertebrate genome and transcriptome sequences that span all major lineages across the vertebrate Tree of Life, including several recently sequenced “ancient fishes” that include bowfin (*Amia calva*), spotted gar (*Lepisosteus oculatus*), and sterlet (*Acipenser rutherens*) that fill in critical phylogenetic nodes absent in earlier macroevolutionary investigations of TLRs (Braasch et al. 2016; Du et al. 2020; Thompson et al. 2021). Using these data, we deploy a series of comparative phylogenetic analyses to estimate the evolutionary history of TLRs across early vertebrate divergences. Our novel *Polypterus* transcriptomes reveal new TLR sequences not identified in existing public genomes that, when placed into a time-calibrated evolutionary framework, aid in identifying a previously unrecognized pulse of TLR diversification. We further investigate the mode of subsequent lineage-specific gains and losses, including consideration of possible life history effects on TLR copy number in teleosts, and reconcile exceptionally high rates of annotation errors from existing genomes with our phylogenetic analyses. Collectively, these results illuminate a novel pattern of innate immune diversification in the wake of the origin of the adaptive immune system while providing a critical reference for the nomenclature of one the most studied innate-immune receptor families in vertebrates.

## Methods

### Data acquisition

We selected 68 representative vertebrate genomes that collectively span all major lineages of the vertebrate Tree of Life : Cyclostoma (2); Chondrichtyes (3); Actinopterygii (32); Actinista (1); Amphibia (2); Lepidosaur (5); Archosaur (7); and Mammalia (18). Genomes for these lineages were downloaded from either NCBI or Ensembl (**Supplementary Table S1**). Our expanded sampling of non-teleost fishes that captures the entire backbone of early diverging Actinopterygiian phylogeny differs from previous investigations of vertebrate TLR receptors (Roach et al. 2005; Alcaide and Edwards 2011; Nie et al. 2018; Liu et al. 2020). This expanded sampling includes Polypteridae, Acipenseriformes (sturgeon and paddlefish), and Holostei (gars and bowfin), thereby allowing us to not only discern the impact of the teleost genome duplication on TLR diversification, but also prevent erroneous assignment of TLR diversity as “teleost-specific” due to taxon sampling that is phylogenetically biased towards only teleosts.

Two novel *Polypterus senegalus* transcriptomes were assembled for use in this study (**Supplementary Text 1** and **Supplementary Table S2**). Polypteriformes are the earliest diverging ray-finned fishes. The early divergence of this lineage offers critical context concerning the early evolution of innate immune receptors and other gene families for half of all living vertebrates. We additionally supplemented our two *P. senegalus* transcriptomes with *Polypterus endlicheri* (NCBI SRA SRX3153328)*, Erpetoichtys calabaricus* (NCBI SRA SRX1661497) (Hughes et al. 2018b), and *Acipenser ruthenus* (NCBI BioProject PRJNA635364) to have more complete representation of early ray finned fishes.

Predicted coding regions within transcriptomic data were identified using transdecoder (https://github.com/TransDecoder/). TLR sequences were identified from genomes and transcriptomes using HMMER (Eddy 2009) with an e-value threshold of 10 x e^-6^. Training profiles for HMMer were built using a fasta alignment of known TLR toll-interleukin-1 receptor (TIR) domains from humans and representative vertebrates (Thompson et al. 2021). TIR domains were chosen for queries as these regions have been shown to be more well conserved and less variable than other TLR regions (Roach et al. 2005; Wu et al. 2018). Moreover, TIR domains exhibit higher levels of sequence conservation than TLR ectodomains which are characterized by leucine rich repeats that are found in over 6000 other proteins on the Pfam database (Matsushima et al. 2007). Putative TLRs were aligned to the reference sequence dataset for manual inspection following phylogenetic verification (see next section). All protein sequences were aligned using MAFFT (Katoh et al. 2002) and inspected via Aliview (Larsson 2014) or Geneious Prime (Biomatters, Inc., San Diego, CA).

### Verification of TLR sequences

Putative TLR sequences were assessed for presence of TLR specific protein domains using HMMER’s hmmscan function against the Pfam database (Mistry et al. 2021). The ultraconserved TIR domain specifically was used to differentiate TLRs from other proteins that possess leucine rich repeats (LRRs). These results were parsed using custom scripts to quantify aspects of the TLR variable ectodomain containing the leucine rich repeats as well as the TIR domain for qualities including domain length and number of domains. Because of the high level of LRR variability within the ectodomain (e.g. sequence variability and number of LRR domains), the TIR domains of only TLR protein sequences were used for subsequent phylogenetic analyses.

### Phylogenetic Analyses of TLRs

TLR structure and the interaction of different domains with PAMPs or signaling pathways has created a scenario where the PAMP-interacting ectodomain evolves at a much faster rate than the signaling TIR domain (Beutler and Rehli 2002; Mikami et al. 2012) and across large evolutionary distances, the LRR domains become unalignable (Roach et al. 2005). TLR identity and evolutionary history was assessed using maximum likelihood based phylogenetic inference of the TIR domains. To find the best fit models of amino acid substitution and infer a maximum likelihood phylogeny of all TIR sequences, alignments generated in MAFFT (Katoh et al. 2002) were analyzed in IQ-TREE2 (Minh et al. 2020) with the candidate pool of substitution rates spanning all common amino acid exchange rate matrices (JTT, WAG, etc). These models also include protein mixture models like empirical profile mixture models, as well as parameters to accommodate among-site rate variation (discrete gamma or free rate model). The best fit model for each alignment was selected using Akaike information criterion with node support assessed via 1,000 ultrafast bootstrap replicates and the resulting tree is available on Zenodo (doi:10.5281/zenodo.7787075).

### Nomenclature for TLRs

Given that TLR sequence names have been reported from a variety of vertebrates inconsistently, we use the gene names from humans (Tweedie et al. 2021), zebrafish (*Danio rerio*) (Traver and Yoder 2020; Bradford et al. 2022), and bowfin (Thompson et al. 2021) as a starting reference for gene nomenclature. We then name TLR sequences relative to these sequences and the reference phylogeny. For example if a sequence named TLR2 in *Anolis* was found to belong in a clade of TLR18, this sequence becomes TLR18 for *Anolis*. Throughout this manuscript, we refer to genera when needing to differentiate between TLRs from different lineages (e.g. *Anolis* TLR18 or *Ursus* TLR10). As we included three species of *Polypterus* and two species of *Salmo* in our taxon-sampling, differentiation in these lineages is at the level of species. Novel TLR sequences identified during the course of this study that are not members of reference clades are named consecutively *TLRnovel1*, *TLRnovel2*, etc.

### Gene Loss and Gain

A time-calibrated tree topology for vertebrates in this study was taken from timetree.org and supplemented with a time-calibrated phylogeny of Acanthomorpha (Ghezelayagh et al. 2022). We applied stochastic character mapping (Bollback 2006) to the present diversity of TLRs to reconstruct the history of gains and losses for each respective TLR. For each TLR, we quantified the probability of ancestral states for each node in the phylogeny based on 1,000 stochastic character maps under the best fit model of evolution selected using Akaike Information Criterion (Akaike 1973). All analyses were conducted using the phytools package in R (Revell 2012). Using the identified origin of each TLR, we then plotted the accumulation of vertebrate TLRs over time. This approach mirrors the commonly used lineage through time plot for assessing the accumulation of taxonomic diversity over time (Nee et al. 1992) that are widely used in evolutionary biology to analyze temporal trends in lineage diversification (Near et al. 2012; Silva et al. 2019; Benton et al. 2022). Analogously, we used this approach to assess whether there were pulses of TLR diversification over the early evolutionary history of vertebrates. In addition to gene gains, we further evaluated the history of gene losses by tracking where in each phylogeny the most probable loss events occurred, generating a stream plot of this history using the ggplot2 package in R (Wickham 2011).

## Results and Discussion

### Identification of twenty-two definitive vertebrate TLR clades

We identified a total of 871 TLR sequences from genomes or transcriptomes of 70 representative vertebrate genera (**Fig. 1**, **Supplementary Fig. S1, Supplementary Tables S1** and **S3**) including two newly generated *Polypterus senegalus* transcriptomes generated for this study (**Supplementary Text 1** and **Supplementary Table S2**). Nearly all of these sequences cluster into 22 TLR clades. With the exception of one TLR-like group of polypteriform TLRs, these 22 clades are strongly supported to belong in six previously described subfamilies by our phylogenetic analyses: TLR1 (bootstrap support [BSS] = 96), TLR3 (BSS = 97), TLR4 (BSS = 99), TLR5 (BSS = 97), TLR7 (BSS = 83), and TLR11 (BSS = 99) (Roach et al. 2005). Within each of the TLR subfamilies, the phylogenetic relationships between TLRs of different taxa largely reflect the evolutionary history of vertebrates. Across this evolutionary history, our results reveal a complex history of TLR gains and losses (**Fig. 1**) that is detailed for each TLR subfamily in **Supplementary Texts** and visualized in **Supplementary Figs. S2-S6**.

**Figure 1.**
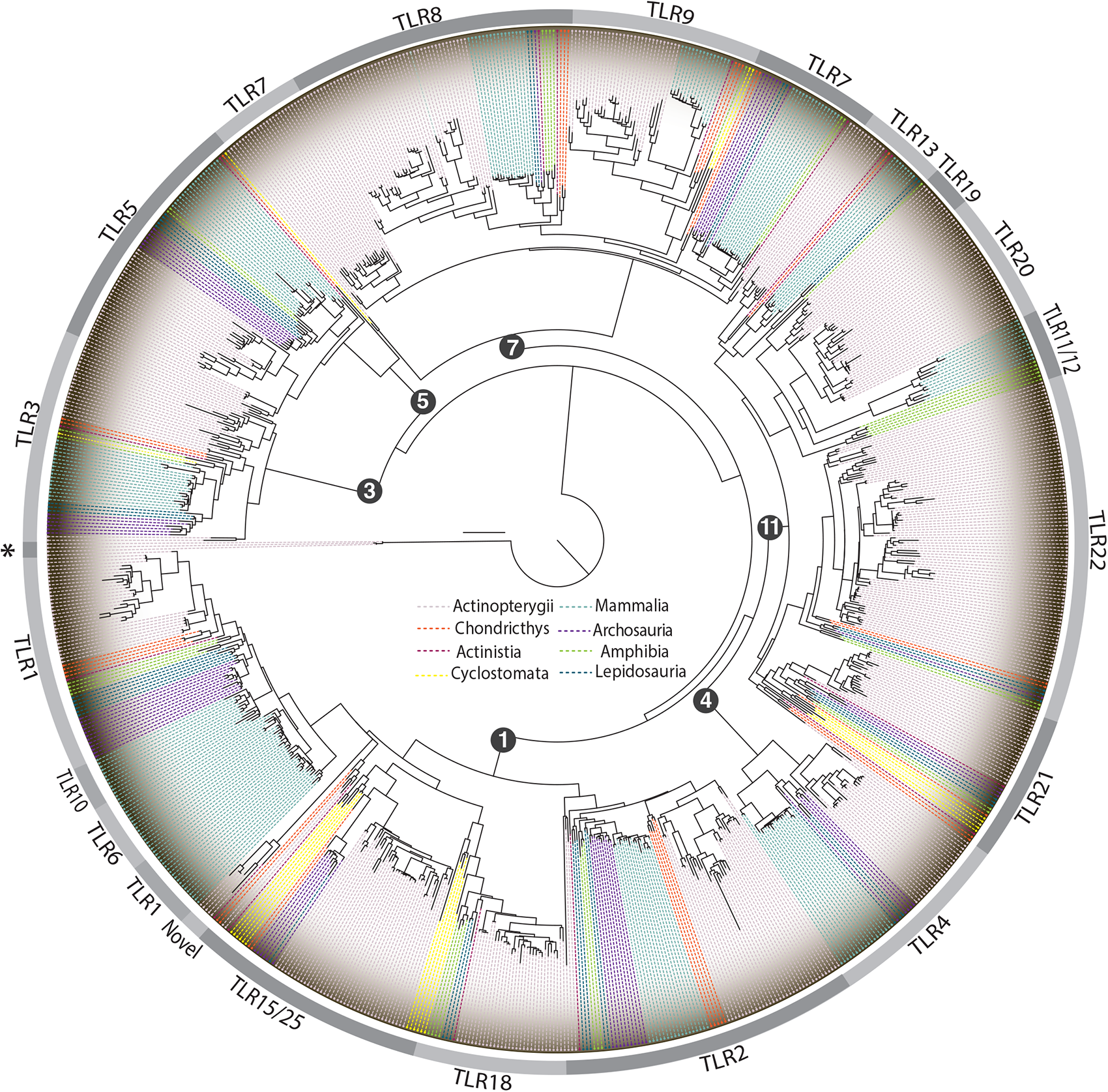
Comparison of representative vertebrate TLRs. Phylogenetic tree of Toll-like receptor TIR sequences identified in 70 genera, from cyclostomes to mammals, representing the breadth of vertebrate taxa with the outer circle designating family boundaries. Sequences are designated as Novel or TLR-like if they do not cluster within other TLR group or subfamily designations, or only appear in one species or lineage (see main text). A group of TLR-like sequences were found in Polypteriformes and used as an outgroup designated by an asterisk. The nomenclature used for major nodes does not always reflect the identity of all sequences in that node. For example, TLR11, TLR12 and a few TLRnovel2 sequences from Leishan spiny toad (*Leptobrachium leishanense*) are labeled as TLR11/12 due to space limitations. See supplementary documents for linear phylogeny (**Supplementary Fig. S1**), trees of subclades (**Supplementary Figs. S2-S6)** which depicts clades at a finer resolution, and accession numbers (**Supplementary Table S1**).

Our results are concordant with several findings from previous attempts to resolve the evolutionary history of vertebrate TLRs (Höhne et al. 2021). For example, we also recover strong support (BSS = 100) that TLR6 and TLR10 are mammal-specific TLRs that likely arose following a duplication event (Huang et al. 2011; Mikacenic et al. 2013) (**Fig. 1** and **Supplementary Fig. S2**). However, we also reveal multiple instances in which the current TLR nomenclature does not align with the major clades defined in **Fig. 1** and **Supplementary Fig. S1**. TLR nomenclature across vertebrate lineages is often in flux making it difficult to directly compare across studies that use different nomenclature and small subsets of samples. For example, TLR14 recently has been referred to as “fish-specific” (Liu et al. 2022), however it also has also been described in *Xenopus* (Roach et al. 2005; Ishii et al. 2007a). On public databases, however, these *Xenopus* sequences have been redefined as TLR2. Our strongly supported resolution (BSS = 100) places these *Xenopus* TLR2/TLR14 sequences within an inclusive TLR18 clade **(Supplementary Figs. S1** and **S3**). Further, the TLR14 name originated from *Takifugu rubripes* and zebrafish sequences in 2005 (Roach et al. 2005; Ishii et al. 2007b, a), but these genes have since been renamed TLR18 in both species. This example highlights the confusing, inconsistent nature of TLR nomenclature throughout the field. We additionally identify a TLR12 from *Xenopus,* previously defined as TLR16 (Roach et al. 2005), that does not fall within any specific clade. This sequence is strongly supported (BSS = 90) as belonging to a clade composed of TLR11, TLR12, TLR19, and TLR20. This divergence of *Xenopus* TLR12 from other TLRs is consistent with results from smaller scale studies on TLRs noting high sequence dissimilarity (Temperley et al. 2008; Roach et al. 2005). The specific resolution of this sequence relative to these strongly supported clades remains elusive (BSS <70; **Supplementary Fig. S6)**, and future studies targeting amphibian TLRs are needed to place this sequence into its evolutionary context.

### Novel TLRs and TLR-like Sequences

The majority of TLR sequences analyzed in this study could be classified into specific subfamily groups (**Fig. 1** and **Supplementary Figs. S1-S6**). However, 13 sequences either formed unique clades within one of the six major subfamilies, or were not resolved within these well supported subfamilies. Instead, these sequences were resolved to reflect three novel TLR lineages. The first of these novel TLR clades is a strongly supported (BSS = 100) novel TLR that is unique to bowfin and spotted gar belonging to the TLR7 subfamily (TLRnovel1; [TLR-HS in (Thompson et al. 2021)]; sequence identifiers 4 and 92 in **Supplementary Table S1**. These two lineages comprise Holostei, one of the earliest diverging lineages of ray-finned fishes (Near et al. 2014; Giles et al. 2017; Dornburg and Near 2021). Our finding of this novel TLR is consistent with an analysis using a more restrictive taxonomic sampling of vertebrates (Thompson et al. 2021), supporting the hypothesis that Holosteans encode a unique TLR (**Supplementary Fig. S5**).

As noted above, five sequences from the Leishan spiny toad (*Leptobrachium leishanense*) were also classified as novel (TLRnovel2: sequence identifiers 830–834 in **Supplementary Table S1**). These sequences fall within the TLR1 subfamily, but were not resolved within other TLR1 clades. However, these sequences are strongly supported (BSS =100) to form the sister lineage to a well supported clade (BSS=90) that comprises TLR19, TLR20, TLR11, TLR12, and *Xenopus* TLR12 **(Supplementary Fig. S6).** Our finding is consistent with work by Babik et al (2014), who noted that the two amphibian TLR12 sequences in their study were highly divergent from other vertebrate sequences and cautioned that the classification of amphibian TLR12 should be treated with caution. Likewise, a recent survey of amphibian TLRs (Zhang et al. 2022) again recovered a clade of amphibian TLR12 sequences with branch lengths indicating high sequence divergence. A more detailed study on the divergence and function of these receptors in amphibians is warranted.

Our third clade of novel TLRs (TLRnovel3; TLR1 subfamily) represents an assemblage that is largely comprised of sequences from early diverging jawed vertebrates including the spotted gar (sequence identifier 105 in **Supplementary Table S1**), coelacanth (*Latimeria*; sequence identifier 88 in **Supplementary Table S1**), and Polypteriformes (sequence identifiers 120, 126, 143 in **Supplementary Table S1**). Previous studies of coelacanth TLRs revealed a highly divergent sequence whose phylogenetic placement was not strongly supported (Boudinot et al. 2014). This sequence was classified as TLR22-like, as it was resolved outside of a clade of TLR22 sequences that included sharks, amphibians, and ray-finned fishes. Bootstrap support for the relationships among these sequences was also low, likely due to limited taxon sampling (Townsend and Lopez-Giraldez 2010). However, our expanded taxon sampling strongly supports (BSS = 96) the resolution of these sequences as a novel TLR clade within the TLR1 subfamily (**Fig. 1** and **Supplementary Fig. S2**). These sequences are resolved with moderate support (70 < BSS > 90) as belonging to a subclade within the TLR1 subfamily containing the well supported TLR25-like (BSS = 100), TLR27 (BSS = 100), and TLR25 clades (BSS = 100). Together with TLR25-like and TLR27, this clade likely represents the vestiges of a round of paralog diversification that occurred in the early evolutionary history of jawed vertebrates, with the notable exception of our finding a sequence within the wrasse (*Labrus bergylta*) genome that is included in this clade **(Supplementary Fig. S2**; sequence identifier 465 in **Supplementary Table S1**). If true, such a finding would suggest either the possibility of an extremely high number of independent gene losses across the history of the approximately 30,000 species of teleost fishes, or that this TLR is more widespread in teleosts than currently recognized. It is also possible that this sequence represents contamination of unknown origin. Studies focusing on the diversification of the ray-finned fish TLR repertoire are needed to disentangle these hypotheses.

In addition to the three novel TLR lineages we also identified a clade of TLR-like sequences unique to Polypteriformes (**Fig. 1**). We resolve a well supported (BSS = 100) clade that contains three sequences from *Polypterus bichir, P. endlicheri*, and *Erpetoichthys calabaricus* (sequence identifiers 121, 137 and 144 in **Supplementary Table S1**). Whether these are true TLRs remains unclear. On the one hand, these sequences are phylogenetically distant from all other TLR sequences in our analysis, and possess the highest degree of sequence divergence. On the other hand, these sequences all possess a TIR domain. Given the ambiguity surrounding the identity of these sequences, we consider them to be TLR-like (**Fig. 1** and **Supplementary Fig. S4)**.

### TLR diversity is conserved but highly asymmetric between lineages

Mapping the diversity of TLRs across vertebrates reveals five out of six major TLR subfamilies are present across all vertebrates, with the the single-gene TLR4 subfamily arising prior to the most recent common ancestor of sarcopterygians and actinopterygians (**Fig. 2**). This result is consistent with the observation that families of vertebrate TLRs are evolutionary conserved (Liu et al. 2020). However, our results also reveal a striking asymmetry in the numbers of paralogs present between species (**Fig. 2**). For example, contrasting the diversity of TLR receptors between major vertebrate clades reveals an imbalance in TLR receptor diversity between jawed (gnathostome) and jawless (cyclostome) vertebrates, with jawed vertebrates generally having twice as many named TLRs as cyclostomes (**Fig. 2**). Similar patterns of asymmetry are also found within major clades. In the TLR11 family, paralog expansions were common in several teleosts with multiple copies of TLR20 in various species as well as large expansions of TLR22 in others such as mudskippers, *Periophthalmus*, that are likely correlated with lineage expansions to novel environments (You et al. 2014; Qiu et al. 2019).

**Figure 2.**
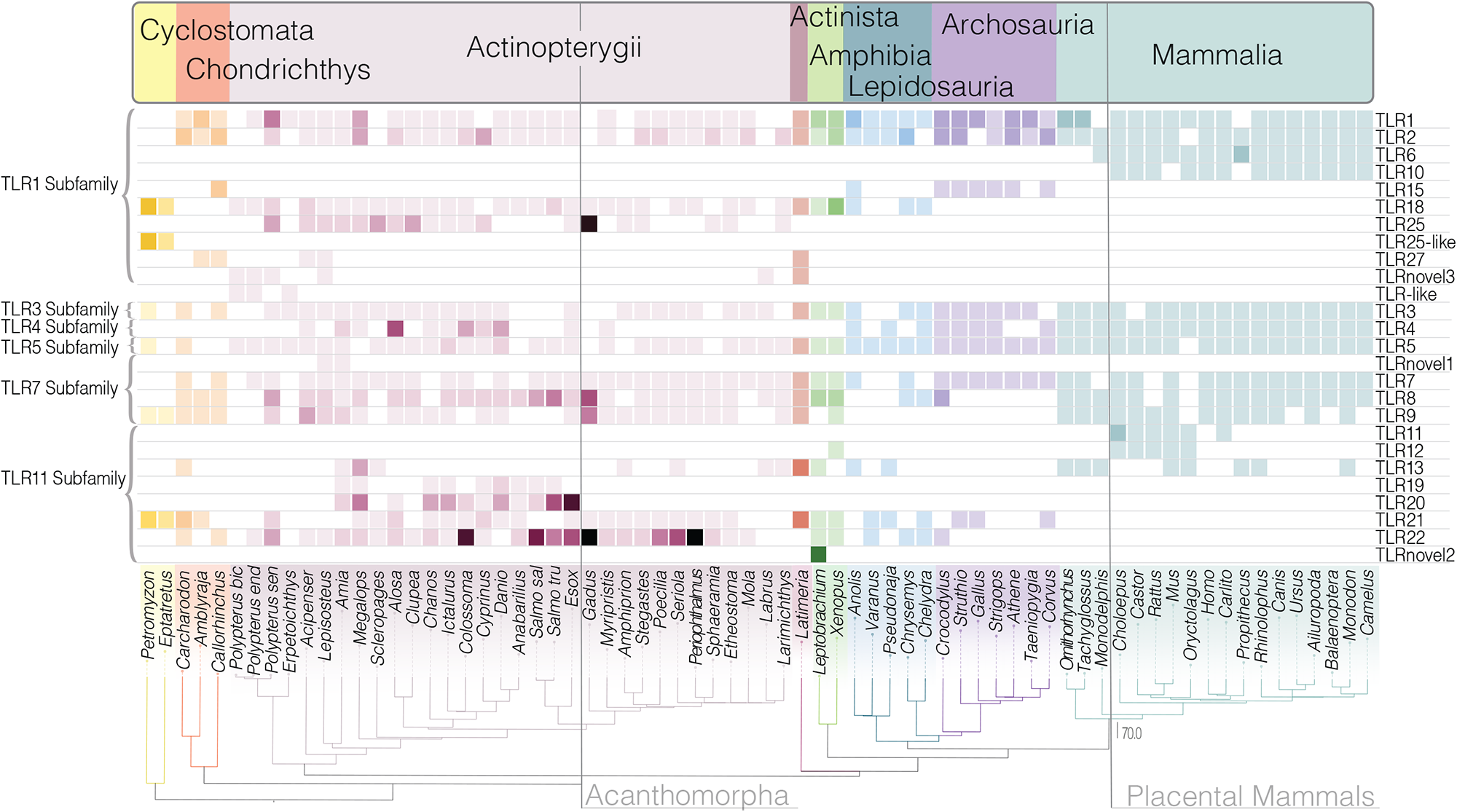
TLR copy number variation. Heatmap of all TLR sequences identified in 70 representative vertebrate genera and the variation present among them. Number of TLRs found is represented by color gradients of light (1) to dark (6+). White squares indicate no TLR sequence was identified while dark/black squares indicate 7 or more TLR sequences were found. Extreme cases include: *Colosomma* TLR22 (7); *Gadus* TLR25 (9), TLR22 (11); *Esox* TLR20 (7); *Periophthalmus* TLR22 (10).

In general, paralog diversification for named TLRs appears most pronounced in ray-finned fishes. Changes in the number of genes between ray-finned fishes and other vertebrates have often been attributed to the teleost genome duplication event; an extra round of genome duplication that occurred prior to the most recent common ancestor of all teleosts (Opazo et al. 2013; Zhang et al. 2018; Parey et al. 2022). However, our results align with a growing consensus that TLR diversification reflects a more complex evolutionary history (Solbakken et al. 2017). We reveal levels of paralog diversity for individual TLRs in non-teleost ray-finned fishes that diverged prior to the teleost genome duplication event including holosteans and Polypteriformes that are on par with, if not exceeding levels, of paralog diversity in teleosts (**Fig. 2**). Additionally, we reveal that the diversity of paralogs is not distributed evenly between teleost species. *Gadus morhua*, for example, has expanded TLR25 and TLR22. As this species has also lost regions of the MHC Class II pathway, it may be under selective pressure to maintain a wider diversity of innate immune resources (Solbakken et al. 2016). These examples illustrate a more general trend of lineage specific paralog diversification that has been found to be a hallmark of many vertebrate gene families (Nei and Rooney 2005; Niimura and Nei 2007; Moriya-Ito et al. 2018; Dornburg et al. 2022). Given that paralog expansions are often linked to aspects of organismal life history and ecology (Moriya-Ito et al. 2018; Corrochano 2019), our analyses can serve as a basis to contextualize the diversity of TLRs in additional taxa. With the decreased cost of genome sequencing, this framework for comparative investigations can facilitate investigations integrating immunological sequence data with other ecological, behavioral, or life history traits for specific lineages, as well as studies attempting to understand the evolution of TLR function.

### Vertebrate TLRs exhibits marked pulses of gene gains and gene losses

Gene families across the Metazoan Tree of Life often depict a signature of paralog diversification and gene loss, with deviations from a continual process often coinciding with pulses of functional innovation (Ota and Nei 1994; Hughes et al. 2018a; Fernández and Gabaldón 2020; Wcisel et al. 2023). Lineage specific expansions have been invoked to explain the evolutionary blooms of vertebrate TLRs based on the stark contrast in TLR diversity between non-vertebrate and vertebrate lineages (Buckley and Rast 2015; Nie et al. 2018; Liu et al. 2020), and the asymmetry in TLR diversity between representative mammals and teleost fish (Sundaram et al. 2012; Solbakken et al. 2016; Khan et al. 2019; Smith et al. 2019). However, our results reveal the potential for a previously undescribed burst of TLR diversification. Our analyses indicate TLR gains are mainly restricted to ancient evolutionary events, with the largest pulse of novel TLR origination coinciding with the origin of gnathostomes (jawed vertebrates; **Fig. 3a** **and** **Fig 3b**). This pulse of diversification is consistent with a hypothesis that the origin of adaptive immunity may have promoted a bout of functional diversification in innate immune receptors. Conversely, it is also possible that the stark contrast in TLR diversity between cyclostomes (jawless vertebrates) and gnathostomes represent a cyclostome specific pulse of TLR gene loss. Further studies of the diversification dynamics of other innate immune receptors shared across vertebrates will be needed to provide evidence for or against these two competing resolutions. Regardless, our analyses support the hypothesis that vertebrate innate immune receptors experienced a burst of diversification during the rise of adaptive immunity.

**Figure 3.**
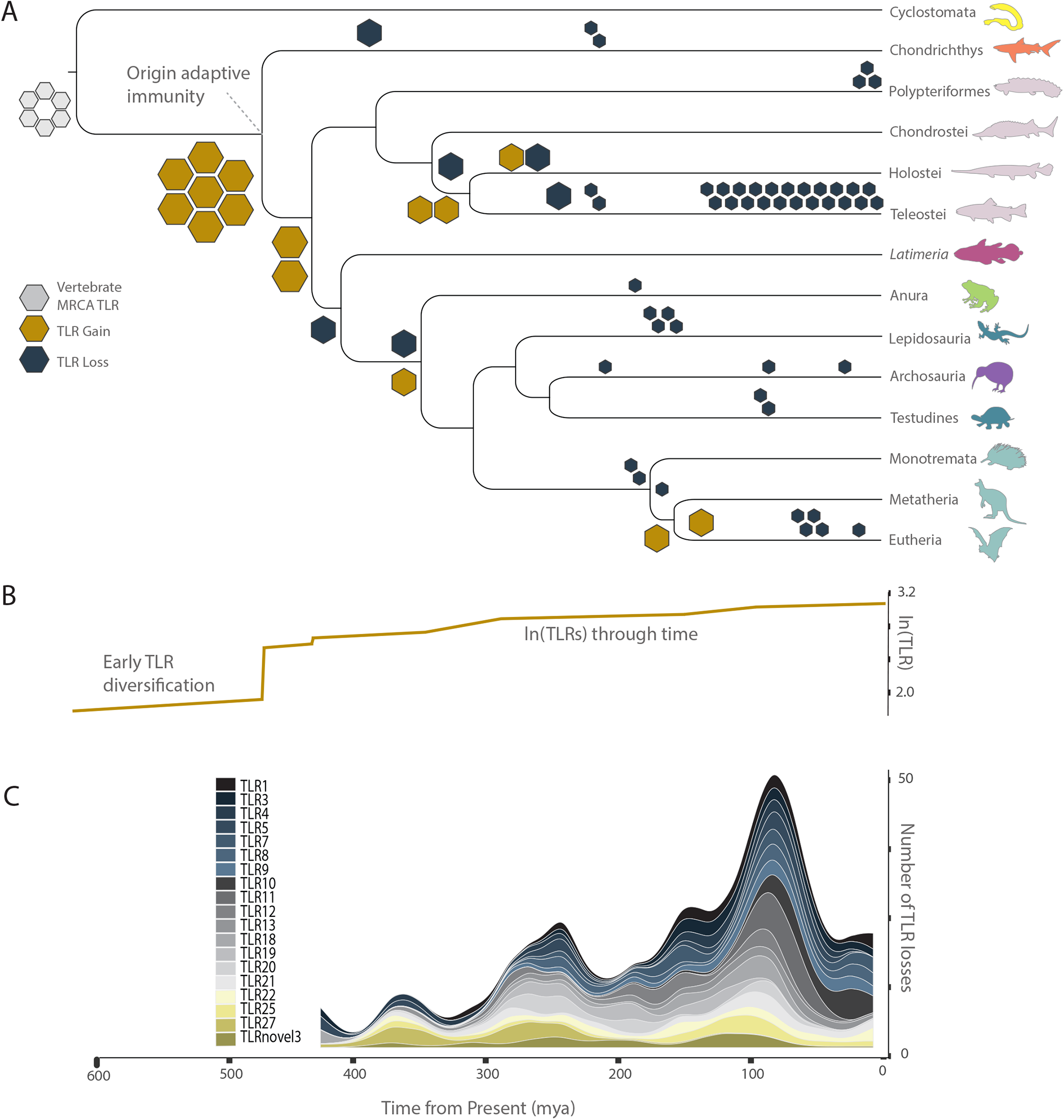
TLR gains and losses. (**a**) Model for loss and gain of vertebrate toll-like receptor (TLR) genes after the rise of the adaptive immune system. The location and number of hexagons are representative of the number of, and the time in which, TLRs were gained (gold hexagons) or lost (dark blue hexagons). (**b**) Natural log of TLR diversification over time. (c) The number of TLR genes lost across six TLR subfamilies over the past 400 million years is presented graphically via a stream graph plot. Hexagons in **a** are scaled to fit within the confines of the phylogeny and the time scale (bottom) is for all panels.

The potential early burst of diversification found by our analyses marked the most substantial increase of vertebrate TLRs in a single time period. However, several major groups of vertebrate TLRs have origins post-dating this event. For example, TLR originations also occur prior to the most recent common ancestor (MRCA) of sarcopterygians and actinopterygians, before the divergence of Holostei and Teleostei, and twice during mammalian evolution — once preceding the split between metatherians and eutherians and then again during eutherian evolution (**Fig. 3a**). Our results reveal how these, along with other lineage specific diversification events, have given rise to a general pattern of how groups of TLRs are distributed among vertebrates. Cyclostomes possess TLR3, TLR5, TLR9, TLR18, TLR21, and TLR25-like with varying numbers of sequences between the two representative genera (**Fig. 2**). Chondrycthian species build upon this repertoire, possessing TLR1, TLR2, TLR3, TLR5, TLR8, TLR9, TLR13, TLR21, TLR22, TLR25, and TLR27 (**Fig. 2**). Actinopterygians were found to present the most diverse repertoire of TLRs with no singular TLR conserved across all genera (**Fig. 2**). This diversity predates the teleost genome duplication event (**Fig. 2**), and places ray-finned fishes in stark contrast to the reduced diversity in archosaurs and mammals.

Our study complements and provides a temporal perspective for previous efforts aimed at characterizing the diversity of TLRs between different groups of vertebrates (Beutler and Rehli 2002; Roach et al. 2005; Liu et al. 2020; Zhang et al. 2022). Our study also reveals an under-appreciated axis of TLR evolutionary history — TLR loss. While it is possible that some perceived losses in our study may be artifacts of genome sequencing, mapping the loss of TLR receptors over time demonstrates distinct pulses of gene loss over vertebrate evolution that are unlikely to occur randomly (**Fig. 3C**). For example, the highest pulse of gene loss occurs in the Late Cretaceous between approximately 100-90 million years ago (**Fig. 3C**). The majority of losses within this time period occur within teleosts (**Fig. 2**), specifically, the largely marine radiation of Acanthomorph (spiny-rayed) fishes (**Fig. 3a**). This period also marks one of the largest carbon perturbations in earth’s history: the mid Cretaceous Oceanic Anoxic Event (OAE2) (Wilson and Norris 2001; Takashima et al. 2011; Li et al. 2022; Matsumoto et al. 2022). During the OAE2, anoxic conditions extended from the surface to depths of over 3000 meters throughout much of the world’s oceans (Wilson and Norris 2001; Takashima et al. 2011). It is possible that this pulse of TLR losses is a consequence of severe population bottlenecks associated with this major extinction in an ancestral acanthomorph fish lineage. This scenario is further supported given that the crown ages of many acanthomorph groups such as Acanthuriforms (surgeonfishes, pufferfishes, snappers), Labriformes (Wrasses), and Carangiformes (Jacks, flatfishes, billfishes, sailfishes) hold their origin shortly after this time period (Near et al. 2013; Ghezelayagh et al. 2022). The impact of the OAE2 or other mass extinction events on vertebrate immune diversity remains little explored (Casadevall and Damman 2020), and further investigations of acanthomorph innate immune receptors are needed to assess if the OAE2 had a marked shift on this clade that includes important model species such as sticklebacks, medaka, and Antarctic fishes.

### Phylogenetics and a stable vertebrate-wide TLR nomenclature

TLRs are one of the most investigated gene families in immunological research (Takeda and Akira 2004; Sabroe et al. 2008; Patra et al. 2021; Khanmohammadi and Rezaei 2021), and are routinely included as a representative family of innate immune receptors in comparative studies (You et al. 2014; Huang et al. 2014; Thompson et al. 2021). A decade ago, an online public catalog organized the structural motifs of 2,572 TLR sequences from 121 species (Gong et al. 2010). However, the database is no longer active, and a consistent naming and classification scheme has yet to be put in place for identifying and naming vertebrate TLRs. Based on the number of sequences whose classifications are not supported by phylogenetic analyses, our results suggest that TLR misclassification is more likely as the evolutionary distance from mammals is increased (**Fig. 4**). Nearly half of the amphibian, actinopterygian, and chondrichthyan named TLRs had names in conflict with their resolution within our phylogeny (**Fig. 1, Supplementary Fig. S1** and **Supplementary Table S1**). A possible explanation for elevated levels of discordance between annotations and phylogenetic inferences in these clades could lie in training data and historic training data for gene annotation software. If there was a possible bias towards human or mammalian genome annotations used in training data, this could have led to decreased accuracy of gene predictions. A not mutually exclusive possibility is that the lack of annotations for close relatives of non-model species has also challenged gene annotation. Regardless, if these or other plausible mechanisms largely explain our observations, it is clear that phylogenetic inference offers a means to reconcile conflicting names and could provide a basis for a vertebrate-wide TLR nomenclature.

**Figure 4.**
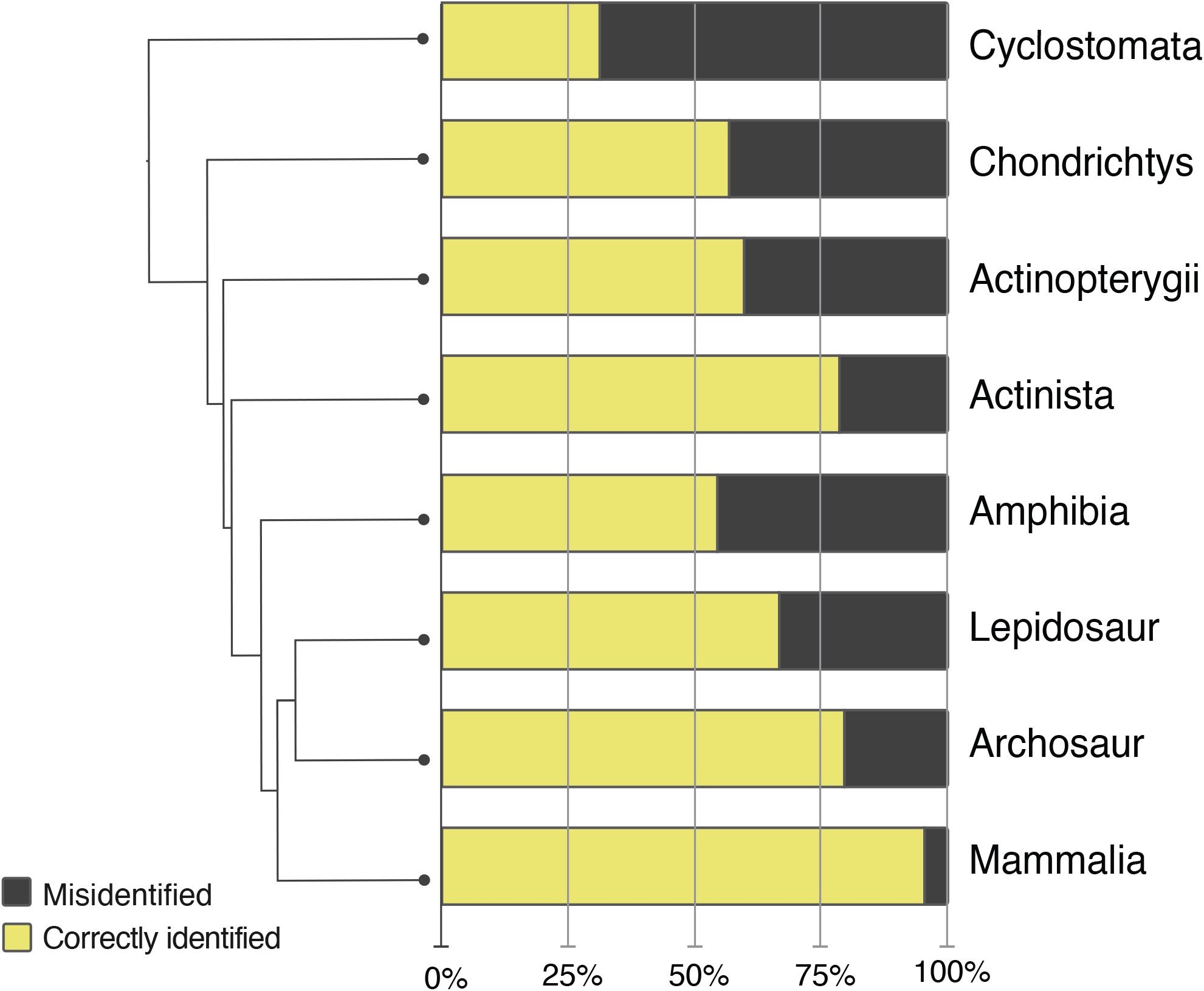
Frequency of misidentified TLRs corresponds to evolutionary distance from mammals. The stacked bar chart indicates correctly identified (yellow) versus misidentified (black) TLRs based on phylogenetic analyses of TIR domains. Taxa are arranged by the phylogenetic relationships of major vertebrate groups, revealing mammals to have the highest level of correctly identified TLRs and cyclostomes having the least. Numbers for calculations are available in supplemental Figure S1, Supplemental Table S1, and the supplemental data files.

Using phylogenetic tree topologies to guide gene nomenclature is not a new idea (Dowse and Soldati 2005; Talbert et al. 2012; Anderson et al. 2016). However, for rapidly evolving domains or genes such as those common in most immune gene families, it is important to consider phylogenetic experimental design choices prior to inference. In the case of TLRs, two approaches have commonly been taken when estimating phylogenetic tree topologies: (1) analysis of the entire gene (Khan et al. 2019; Liu et al. 2020; Zhang et al. 2022) or (2) analysis of only the TIR domain (Ishii et al. 2007a; Boudinot et al. 2014; Thompson et al. 2021). In our study, we restricted our inference to including only the TIR domain, as previous work has indicated that the variable number of and rapidly evolving nature of TLR leucine-rich repeat (LRR) regions renders them unreliable for inference (Ishii et al. 2007a; Mikami et al. 2012). Further, it has become clear that biases in base composition between lineages, such as differences in the leucine content of LRRs between lineages, can mislead inference and even artificially inflate support values (Romiguier et al. 2016; Dornburg et al. 2017; Laumer et al. 2018). We propose that future efforts to reconcile TLR nomenclature carefully consider the underlying data used for tree inference. Such work is outside of the scope of this manuscript as multiple model species such as *Danio, Xenopus, Mus,* and *Homo* each have their own nomenclature committees.

A reconciliation of TLR nomenclature would require a coordinated effort across these and other groups, and could yield a highly useful resource for the immunogenetics community.

## Acknowledgements

We thank Ashley Fletcher for assistance with Polypterus RNA preparations and the NC State University Genomic Sciences Laboratory for running RNA-seq samples. We thank members of the Dornburg and Yoder labs for helpful comments during the early stages of this project.

## Contributions

JAY and AD conceived the project; CN datamined public databases; CN & AD generated code used for analysis; KBC and DJW completed transcriptome analyses; KBC, CN, and AD performed analyses; KBC & AD created graphics; KBC & AD wrote the first draft of the manuscript; JAY and AD supervised the project. All authors read and approved the manuscript.

## Data availability

New transcriptome sequence read data used in this study have been deposited to NCBI under the project accession number PRJNA950356. Custom scripts, alignments, and phylogenetic tree topologies used in this project are available on Zenodo (doi:10.5281/zenodo.7787075).

