## Supplemental Figure 1 for "Ancient Fish Lineages Illuminate Toll-Like Receptor Diversification in Early Vertebrate Evolution"

**Supplementary Figure S1.** Detailed phylogeny with all data from Fig.1 & Supplementary Table S1. which includes bootstrap values, sequence identifiers, sequence names (as designated in NCBI at the time of data mining) for all 870 sequences used in this study. This phylogeny also includes our categorization of 6 TLR subfamilies (black circles with white text), and 28 TLR lineages or groups (red lines, and black bold text). Along with the classical, previously defined TLR groups, we find three novel TLR lineages and a clade of novel TLR-like sequences unique to Polypteriformes. Bootstrap values correspond to circle color. Bootstrap values can be found beside their corresponding circle at each node.
* indicates sequences removed from NCBI

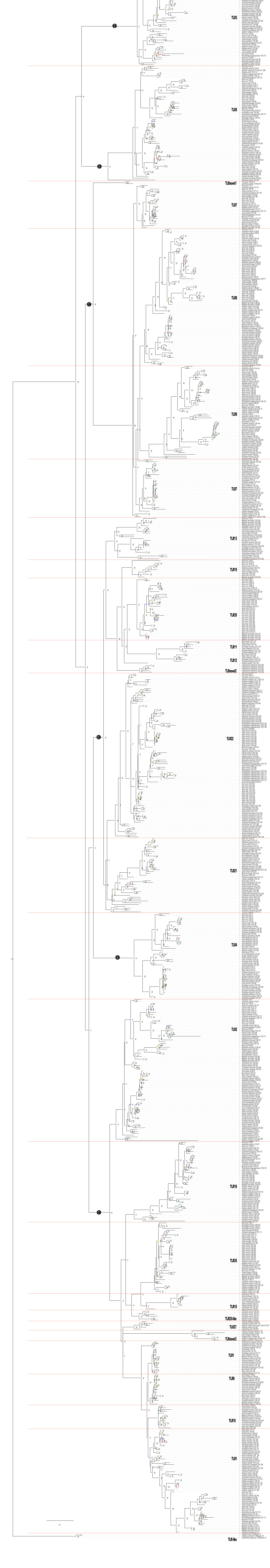
