## Supplemental Materials for "Ancient Fish Lineages Illuminate Toll-Like Receptor Diversification in Early Vertebrate Evolution"

— *Immunogenetics* —

* Corresponding author at:

### *Polypterus senegalus*

*Supplementary Text 1: RNA-seq Methods*

Polypteriformes represent the earliest diverging branch in the ray-finned fish Tree of Life. The early divergence of this lineage offers critical context concerning the early evolution of innate immune receptors and other gene families for half of all living vertebrates. To augment genomic resources for this clade, novel transcriptomes were generated for two *Polypterus senegalus* purchased through the pet trade (individual B0002 and B0003). All research involving live animals was performed in accordance with relevant institutional and national guidelines and regulations, and was approved by the North Carolina State University Institutional Animal Care and Use Committee. The fish were euthanized, and tissues (gill, intestine, and spleen) were dissected. RNA was extracted from each individual tissue and quantified using a NanoDrop 1000 (Thermo Fisher) and Agilent Bioanalyzer. For each individual fish, equal amounts of RNA from each tissue was pooled for sequencing.

mRNA from each tissue was enriched using oligo(dT) beads, rRNA was removed using a Ribo-Zero kit (Epicentre, Madison, WI) and mRNA was randomly fragmented. A cDNA library from pooled RNA from each individual was then prepared for sequencing. Library preparation and sequencing was performed by the North Carolina State University Genomic Sciences Laboratory. Next-gen sequencing (2 × 150 bp paired end reads) was performed on a NextSeq500 instrument (Illumina). Adapter sequences and poor-quality reads were filtered with Trimmomatic v34 (Bolger et al. 2014). The transcriptome was de novo assembled with Trinity v2.11.0 (Grabherr et al. 2011). Benchmarking Universal Single-Copy Orthologs (BUSCO) analysis (Manni et al. 2021) comparing the *P. senegalus* transcriptomes against the Vertebrata and Actinopterygii datasets (**Supplementary Table S2**) revealed ~84% and ~77% coverage, respectively. Raw reads and computationally assembled transcriptome sequences were deposited onto NCBI under the bioproject accession number PRJNA950356.

*Supplementary Table S2: Polypterus senegalus RNA-seq BUSCO results*[1]

| | ***P. senegalus* 0002 vs. vertebrata** | ***P. senegalus* 0002 vs. actinopterygii** | ***P. senegalus* 0003 vs. vertebrata** | ***P. senegalus* 0002 vs. actinopterygii** |
|---|---|---|---|---|
| Complete Total | 84.1% | 76.4% | 84.2% | 77.5% |
| Complete Single | 51.2% | 46.6% | 52.2% | 48.5% |
| Complete Duplicate | 32.3% | 29.8% | 31.9% | 29.0% |
| Fragmented | 7.6% | 4.5% | 6.9% | 4.3% |
| Missing | 8.9% | 19.1% | 9.0% | 18.2% |
| Total BUSCO Groups | 3354 | 3640 | 3354 | 3640 |

[1] Translated *P. senegalus* transcriptomes were assessed for completeness using BUSCO vertebrata and actinopterygii odb10 datasets (Manni et al. 2021)

***Supplementary Table S3: Representative species with scientific and common names***

| Classifications | | | Scientific Name | Common Name |
|---|---|---|---|---|
| **Cyclostomata** | | | *Petromyzon marinus* | Sea lamprey |
| | | | *Eptatretus burgeri* | Hagfish |
| **Chondricthys** | | | *Carcharodon carcharias* | Great white shark |
| | | | *Amblyraja radiata* | Thorny skate |
| | | | *Callorhinchus milii* | Ghost shark |
| **Actinopterygii** | | **Polypteriformes** | *Polypterus bichir* | Nile bichir |
| | | | *Polypterus endlicheri* | Saddled bichir |
| | | | *Polypterus senegalus* | Senegal bichir |
| | | | *Erpetoichthys calabaricus* | Reedfish |
| | | **Acipenseriformes** | *Acipenser ruthenus* | Sterlet |
| | | **Holostei** | *Lepisosteus oculatus* | Spotted gar |
| | | | *Amia calva* | Bowfin |
| | **Teleostei** | **MRCA Elopomorpha and Osteoglossomorpha** | *Megalops cyprinoides* | Indo-Pacific tarpon |
| | | | *Scleropages formosus* | Asian arowana |
| | | **Otocephala** | *Alosa sapidissima* | American shad |
| | | | *Clupea harengus* | Atlantic herring |
| | | | *Chanos chanos* | Milkfish |
| | | | *Ictalurus punctatus* | Channel catfish |
| | | | *Colossoma macropomum* | Tambaqui |
| | | | *Cyprinus carpio* | Common carp |
| | | | *Danio rerio* | Zebrafish |
| | | | *Anabarilius grahami* | Kanglang fish |
| | | **Salmoniformes** | *Salmo salar* | Atlantic salmon |
| | | | *Salmo trutta* | Brown trout |
| | | | *Esox lucius* | Northern pike |
| | | **Acanthomorpha** | *Gadus morhua* | Atlantic cod |
| | | | *Myripristis murdjan* | Pinecone soldierfish |
| | | | *Amphiprion ocellaris* | Clownfish |
| | | | *Stegastes partitus* | Bicolor damselfish |
| | | | *Poecilia formosa* | Amazon molly |
| | | | *Seriola lalandi dorsalis* | Yellow amberjack |
| | | | *Periophthalmus magnuspinnatus* | Mudskipper |
| | | | *Sphaeramia orbicularis* | Orbiculate cardinalfish |
| | | | *Etheostoma spectabile* | Orange throat darter |
| | | | *Mola mola* | Ocean Sunfish |
| | | | *Labrus bergylta* | Ballan wrasse |
| | | | *Larimichthys crocea* | Large yellow croaker |

| | | | | | |
|---|---|---|---|---|---|
| **Sarcopterygii** | | | **Actinistia** | *Latimeria chalumnae* | Coelacanth |
| | **Tetrapoda** | | **Amphibia** | *Leptobrachium leishanense* | Leishan spiny toad |
| | | | | *Xenopus tropicalis* | Western clawed frog |
| | | | **Lepidosauria** | *Anolis carolinensis* | Green anole |
| | | | | *Varanus komodoensis* | Komodo dragon |
| | | | | *Pseudonaja textilis* | Eastern brown snake |
| | | | | *Chrysemys picta bellii* | Painted turtle |
| | | | | *Chelydra serpentina* | Common snapping turtle |
| | | | **Archosauria** | *Crocodylus porosus* | Saltwater crocodile |
| | | | | *Struthio australis* | Southern ostrich |
| | | | | *Gallus gallus* | Chicken |
| | | | | *Strigops habroptila* | Kakapo |
| | | | | *Athene cunicularia* | Burrowing owl |
| | | | | *Taeniopygia guttata* | Zebra finch |
| | | | | *Corvus moneduloides* | New Caledonian crow |
| | | **Mammalia** | **Monotremata** | *Ornithorhynchus anatinus* | Platypus |
| | | | | *Tachyglossus aculeatus* | Short-beaked echidna |
| | | | **Metatheria** | *Monodelphis domestica* | Short-tailed opossum |
| | | | **Eutheria** | *Choloepus didactylus* | Two-toed sloth |
| | | | | *Castor canadensis* | American Beaver |
| | | | | *Rattus rattus* | Black rat |
| | | | | *Mus musculus* | Mouse |
| | | | | *Oryctolagus cuniculus* | European rabbit |
| | | | | *Homo sapiens* | Human |
| | | | | *Carlito syrichta* | Philippine tarsier |
| | | | | *Propithecus coquereli* | Coquerel's sifaka |
| | | | | *Rhinolophus ferrumequinum* | Greater horseshoe bat |
| | | | | *Canis lupus familiaris* | Dog |
| | | | | *Ursus arctos horribilis* | Grizzly bear |
| | | | | *Ailuropoda melanoleuca* | Giant panda |
| | | | | *Balaenoptera musculus* | Blue whale |
| | | | | *Monodon monoceros* | Narwhal |
| | | | | *Camelus dromedarius* | Dromedary camel |

***TLR1 Subfamily (1/2/6/10/15/18/25/27)***

*Supplementary Text 2: TLR1 Subfamily Overview*

The TLR1 subfamily comprises TLR1, TLR2, TLR6, TLR10, TLR15, TLR18, TLR25, and TLR27. Relative to other vertebrates, teleost species generally had the lowest representation of these TLR1 subfamily genes. In general, teleosts possess three or four TLR1 subfamily genes. However, our analyses reveal additional lineage specific losses. For example, we found no TLR1 subfamily genes in *Larimichthys* and only two in *Amphiprion*.

In terms of paralog diversity, multiple copies are uncommon in the TLR1 subfamily. In actinopterygians, TLR2 and TLR25 displayed the most duplications. In placental mammals, having three or four non-duplicated TLR1 subfamily genes was generally the most common state, though some lineages like *Propithecus* only displayed two: TLR2 and TLR6, with two copies of TLR6. Replication was also commonly seen in non-placental mammals, archosaurs, amphibians, and to a lesser degree in lepidosaurs. Duplications of TLR2 were also seen in amphibians, archosaurs, and lepidosaurs (Supplementary Figures S2 & S3).

*Supplementary Text 3: TLR1/6/10*

TLR1 was not observed in cyclostomes and seems to have originated prior to the recent common ancestor of jawed vertebrates. In all vertebrates except for placental mammals, possession of one or two TLR1 sequences was most common with only *Megalops* and *Polypterus* showing evidence of three. In general, most actinopterygians possess a single copy of TLR1 with several genera (*Acipenser*, *Scerophages*, *Cyprinus*, *Gadus*, *Labrus*, and *Larimichthys*) having none. Most sarcopterygians have one or two copies of TLR1. Placental mammals have only a single copy of TLR 1. However, all placental mammals also possess TLR10 and TLR6. TLR6 is also found in *Monodelphis* suggesting an origin following the duplication of TLR1 prior to the most recent common ancestor of metatherians and eutharians. As TLR10 is present in only eutherian mammals, this suggests a second duplication event gave rise to this receptor. TLR1 and are present in nearly every placental mammal genus we surveyed with only two exceptions in *Mus* and *Propithecus*. *Propithecus* does exhibit a duplication of TLR6, which may help to compensate for a lack of both TLR1 and TLR10. In monotremes, TLR6 and TLR10 are not present. Instead TLR1 is duplicated which may suggest a lineage specific paralog diversification event (Supplementary Figure S2).

*Supplementary Text 4: TLR2*

TLR2 is found in all major lineages of jawed vertebrates. In general, most vertebrates possess either one or two copies of TLR2. Notable exceptions include *Cyprinus* and *Megalops* that each encode three copies. In most vertebrates, these gene duplications indicate lineage specific paralog diversification events. For example, duplications were noted in both shark species (*Carcharodon* & *Callorhinchus)* but not in skate (*Ambylata*), suggesting a shark specific duplication event. Likewise, lineage specific duplications were observed within *Chrysemys*, several archosaur species (*Athene, Corvus, Struthio*), and *Xenopus*. However, recent lineage specific duplications do not capture the entire evolutionary history of TLR2 in teleost fishes. Instead, our results suggest that teleost paralogs are an amalgamation of both recent lineage specific duplications, as well as an ancient gene duplication event that gave rise to two TLR clades. This evolutionary history is supported by our phylogenetic analyses that reveal evidence for lineage specific clusters (*Cyprinus, Colossoma, etc)* as well as species that have TLR sequences in both TLR2 clades (*Stegastes, Spharamia*) (Supplementary Figure S3).

*Supplementary Text 5: TLR18*

TLR18 is present in both cyclostome species and is one of only two TLR1 subfamily genes present in Cyclostomata. This suggests that TLR18 may have provided the genetic substrate for the origination of other TLR genes in jawed vertebrates. TLR18 was found to be present in nearly all Actinopterygian genera with only a few exceptions, however it was absent in all chondrichthyans, suggesting a possible loss. It is present in Actinistia, Amphibia, as well as in some lepidosaurs and testudines. However, TLR18 appears to have been lost independently in archosaurs and mammals (Supplementary Figure S3).

*Supplementary Text 6: TLR25 & TLR15*

TLR25 is not monophyletic. Instead, a well-supported clade of TLR15 that includes multiple archosaurs, *Anolis*, and *Callorhinchus* is contained within the TLR25 lineage. Based on our phylogenetic results, this TLR15 clade likely reflects a gene duplication in early jawed vertebrates in which one copy was maintained in sarcopterygians and the other in actinopterygians. In sarcopterygians, TLR15 is single copy. However, in actinopterygians, TLR25 is often duplicated within lineages. The most extreme example occurs in *Gadus*, where we observe 9 copies (Supplementary Figure S2).

*Supplementary Text 7: TLR27*

TLR27 is restricted to a small subset of vertebrates with early divergences that include elasmobranchs (sharks and rays), coelacanth (*Latimeria*), and the earliest divergences in ray-finned fishes (Polypteriformes, Acipenseriformes, and Holostei (Supplementary Figure S2).

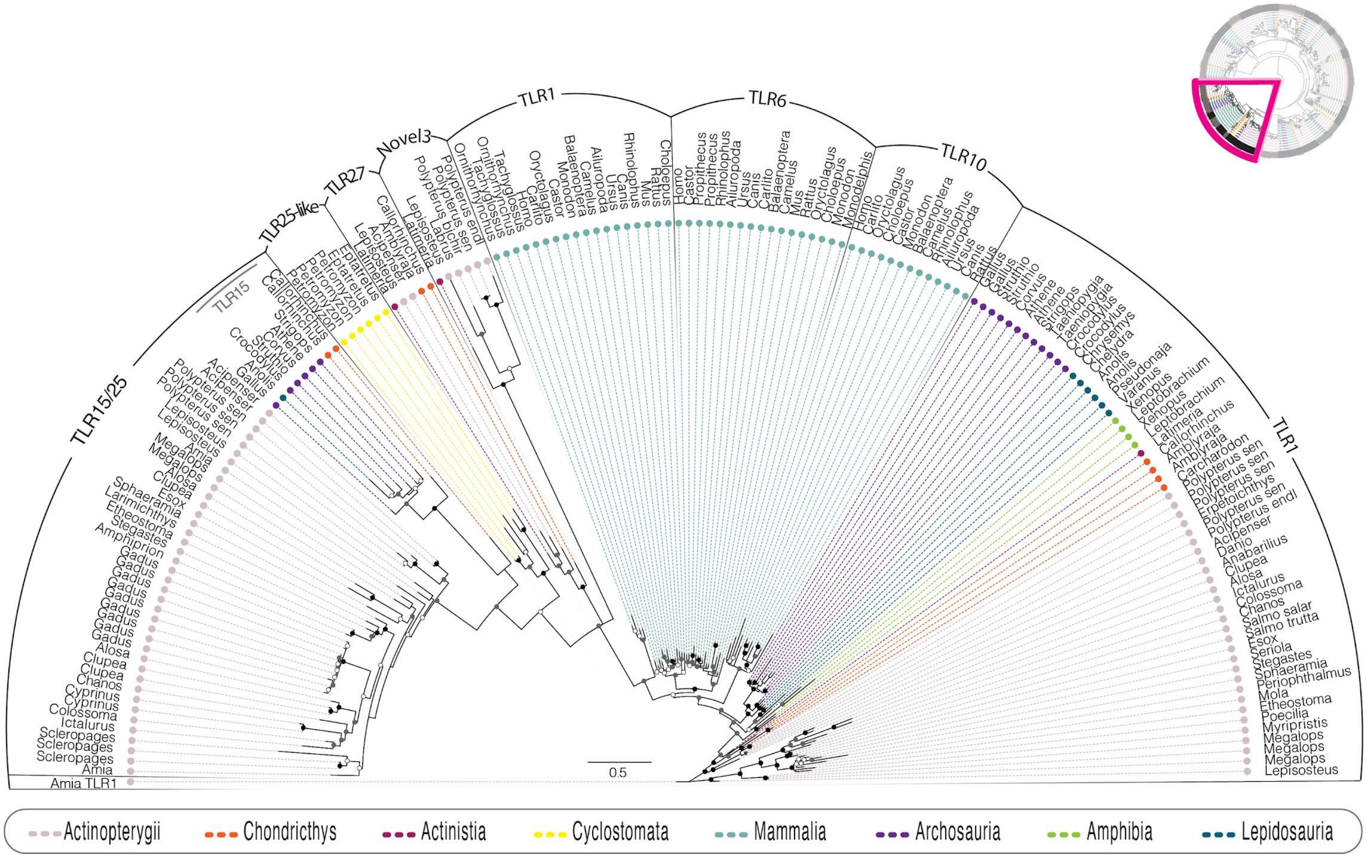

*Supplementary Figure S2. Phylogenetic tree of TLR1 subfamily (partial)*

Subtree of **Fig. 1** which shows the relationship of TLR1 subfamily (partial). Individual sequences are defined by their genus name on the perimeter of the subtree and TLR group designation is indicated by the outermost ring. Position of the subtree within the larger phylogeny (**Fig. 1**) is represented by a pink triangle in the upper right corner. Note the species encoding TLRnovel3 sequences include spotted gar, coelacanth and Polypteriformes. Bootstrap values are represented by black (BSS=100), gray (90 ≤ BSS <100), or white (70 ≤ BSS <90) circles and each class represented by a different color line.

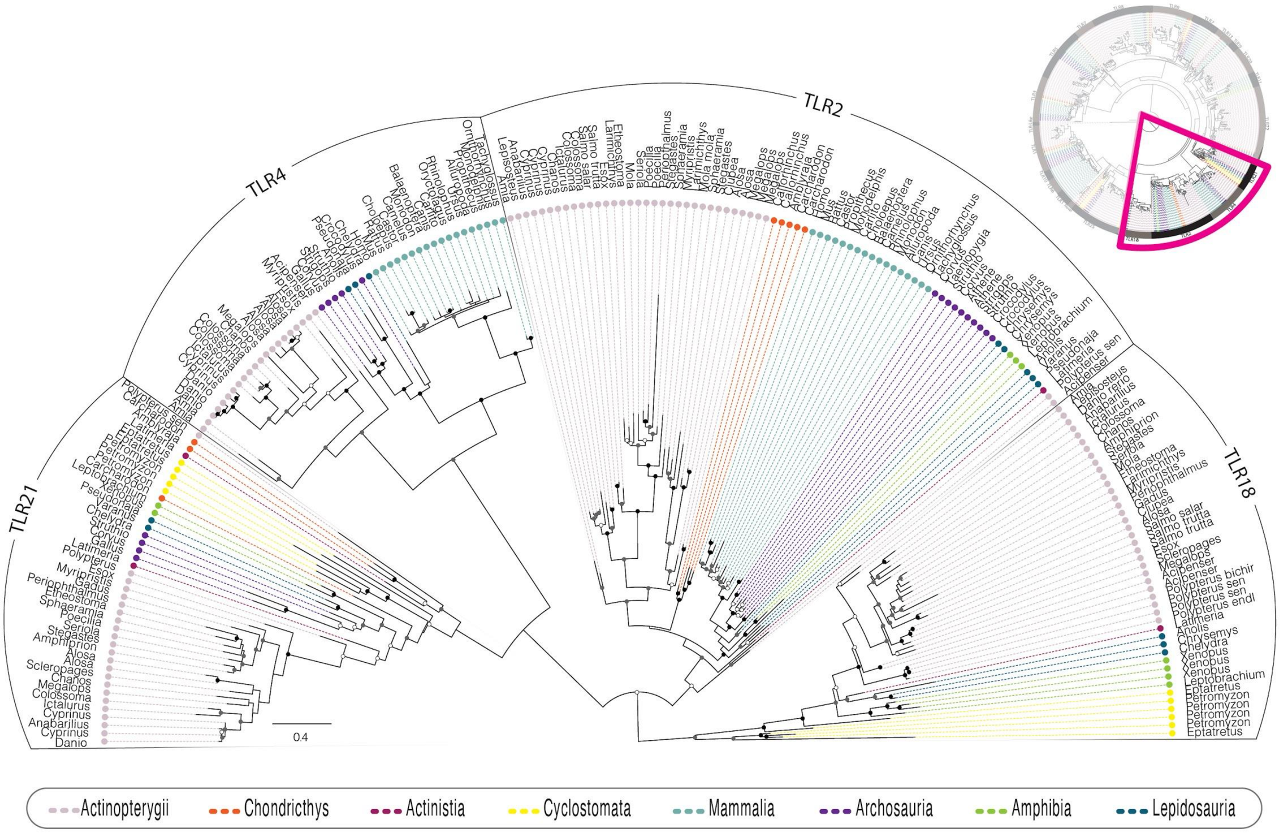

*Supplementary Figure S3. Phylogenetic tree TLR1 (partial), TLR4 and TLR11 (partial) subfamilies*

Subtree of **Fig. 1** which shows the relationship of TLR1, TLR4, and TLR11 subfamilies. Individual sequences are defined by their genus name on the perimeter of the subtree and TLR group designation is indicated by the outermost ring. Position of the subtree within the larger phylogeny (**Fig. 1**) is represented by a pink triangle in the upper right corner. Bootstrap values are represented by black (BSS=100), gray (90 ≤ BSS <100), or white (70 ≤ BSS <90) circles and each class represented by a different color line.

#### ***TLR3, TLR4 and TLR5***

*Supplementary Text 8: TLR3, TLR4, & TLR5 Subfamilies*

TLR3, TLR4 and TLR5 were each resolved as monophyletic with strong support. In sarcopterygians, cyclostomes, and elasmobranchs, these were exclusively single copy. However, some actinopteryians possess multiple copies of TLR4 or TLR5. For TLR4, bowfin possess two copies, and we also observed four otocephalan lineages with 2-5 paralogs per species (*Alosa*, *Colossoma*, *Danio*, *Cyprinus*). All of these except for zebrafish (*Danio rerio*) displayed single copies of TLR5. Only one other actinopterygian species, the channel catfish (*Ictalurus punctatus*), also exhibited a duplication of TLR5. Across all vertebrates, no duplication was observed in TLR3.

*Supplementary Text 9: TLR3*

TLR3 was found in all vertebrate lineages. However, the Petromyozon sequence was well supported as being nested within a clade of amphibian TLR3, suggesting the possibility for error in the publicly available data that warrants investigations outside of the scope of this project. TLR3 also exhibited asymmetry in branch lengths between vertebrate clades, with mammals generally displaying shorter branch lengths compared to other clades (Supplementary Figure S4).

*Supplementary Text 10: TLR4*

Based on our analysis, the TLR4 subfamily likely arose prior to the most recent common ancestor of sarcopterygians and actinopterygians. The TLR4 lineages within these two clades form well-supported reciprocally monophyletic groups. Within actinopterygians, TLR4 appears to have been lost in Polypteriformes, but is present in Acipenseriformes, Holostei, and teleosts. Within teleosts, we find no evidence of TLR4 in any acanthomorph lineages except for the soldierfish, *Myripristis*. Actinistia and amphibians also appear to have lost TLR4 independently. However, TLR4 is present in certain amphibians not included in this study (e.g., *Xenopus laevis*, *Anaxyrus baxteri*, *Bufo bufo*, *Rhinella marina*: Carlson et al, 2022), so our observations likely represent lineage-specific losses in Amphibia. This is similar to our finding that TLR4 is present in most other tetrapod lineages with the exception of tipward lineage specific losses. Of particular note is that TLR4 is seen in every mammalian species. This prevalence likely reflects a potentially critical role in mammalian innate immune response. For example, vertebrates TLR4s possess differing response levels to lipopolysaccharide (LPS). TLR4 is known to bind to LPS in mammals and induce the release of proinflammatory cytokines critical to the immune response (Lu et al. 2008). In contrast, many studied teleosts show nearly no response to LPS injection (Berczi et al. 1966). These differing levels of susceptibility between mammals and teleosts underscore the potential for differing functions of TLR4 across vertebrate taxa (Supplementary Figure S5).

*Supplementary Text 11: TLR5*

TLR5 is found in all major vertebrate lineages as a single copy. Across lineages surveyed, TLR5 was ubiquitous in archosaurs, actinista, lepidosaurs, amphibia and all but one mammal. However, the presence of TLR5 was not uniform within actinopterygians, chondrichthyians, or cyclosomes; suggesting multiple lineage-specific losses.

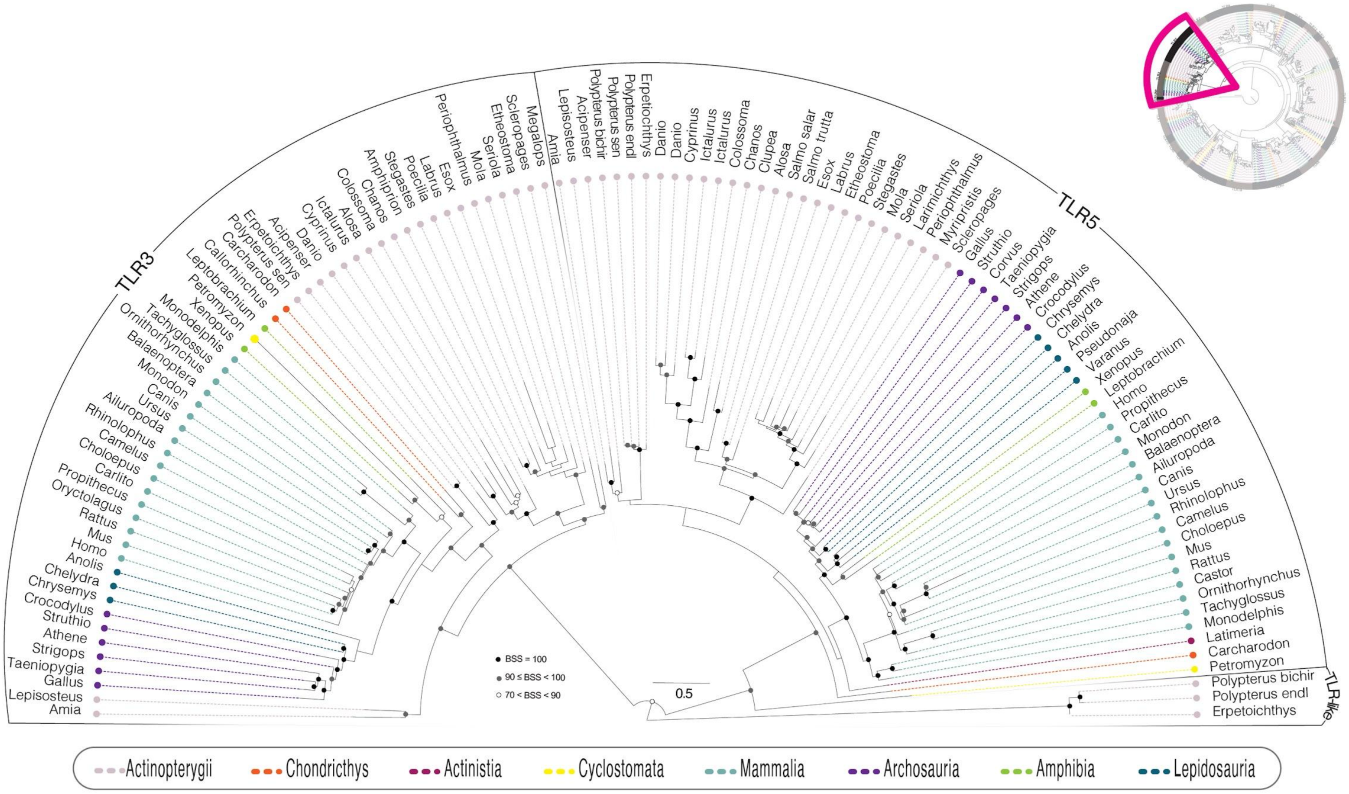

*Supplementary Figure S4. Phylogenetic tree of TLR3 and TLR5 subfamilies*

Subtree of **Fig. 1** which shows the relationship of TLR3 and TLR5 subfamilies and TLR like sequences. Individual sequences are defined by their genus name on the perimeter of the subtree and TLR group designation is indicated by the outermost ring. The position of the subtree within the larger phylogeny (**Fig. 1**) is represented by a pink triangle in the upper right corner. Note the polypteriform-specific clade of TLR-like sequences. Bootstrap values are represented by black (BSS=100), gray (90 ≤ BSS <100), or white (70 ≤ BSS <90) circles and each class represented by a different color line.

#### *TLR7 subfamily (7/8/9)*

*Supplementary Text 12: TLR7*

Our analyses do not resolve TLR7 as monophyletic. However, this lack of resolution is not well supported and reflects a lack of phylogenetic information rather than non-monophyly. We do resolve four strongly supported clades whose relation one another remains undefined: (1) Teleostei + Holostei; (2) *Polypterus* & *Acipenser*; (3) Chondrichthyes; and (4) Sarcopterygii. Given the distribution of vertebrate TLR7, this receptor seems to have arisen prior to the most recent common ancestor of jawed vertebrates. A single copy of TLR7 is virtually ubiquitous across vertebrates, with no additional paralogs observed for any species in this study.

*Supplementary Text 13: TLR8*

TLR8 is strongly supported as monophyletic by our analyses and arose in the most recent common ancestor of jawed vertebrates. Like TLR7, TL8 is found across most vertebrates in our studies with few exceptions due to lineage-specific losses that include a potential loss of TLR8 in the most recent common ancestor in living birds. Lineage-specific paralogs were observed in both sarcopterygians and actinopterygians. However, the extent of paralog diversity varies between these clades. Within sarcopterygians, lineage-specific paralogs were restricted to our representative crocodilian and Amphibia.

In contrast lineage-specific paralogs were relatively common in most Actinopterygian genera except for acanthomorphs, suggesting the possibility of shifts in the rate of TLR8 paralog diversification across vertebrates.

*Supplementary Text 14: TLR9*

TLR9 was found in all vertebrate lineages, including cyclostomes, indicating an origin prior to the most recent common ancestor of all vertebrates. In most vertebrates, TLR9 is a single copy gene. However, there are notable exceptions to this pattern in actinopterygians. First, non-teleosts (Polypterus, sturgeon, bowfin, and gar) all possess two copies of TLR9. In contrast, most teleosts possess only a single copy, suggesting a loss of this receptor following the teleost genome duplication event. Second, there is a major expansion of TLR9 in *Gadus*. Third, there are numerous lineage-specific losses of TLR9 — for example, both *Salmo* species investigated lack TLR9. TLR9 was also not found in any archosaur, lepidosaur, or testudine lineages. We also observed a dramatic reduction in TLR9 branch lengths within placental mammals relative to any other vertebrates, indicating a potential shift in the rate of molecular evolution. Such a slowdown may be explained by the observation that TLR9 may be under stronger stabilizing selection (Jiang et al. 2007). Future tests of TLR9 function in other vertebrates are needed to discern the explanatory power of such a hypothesis.

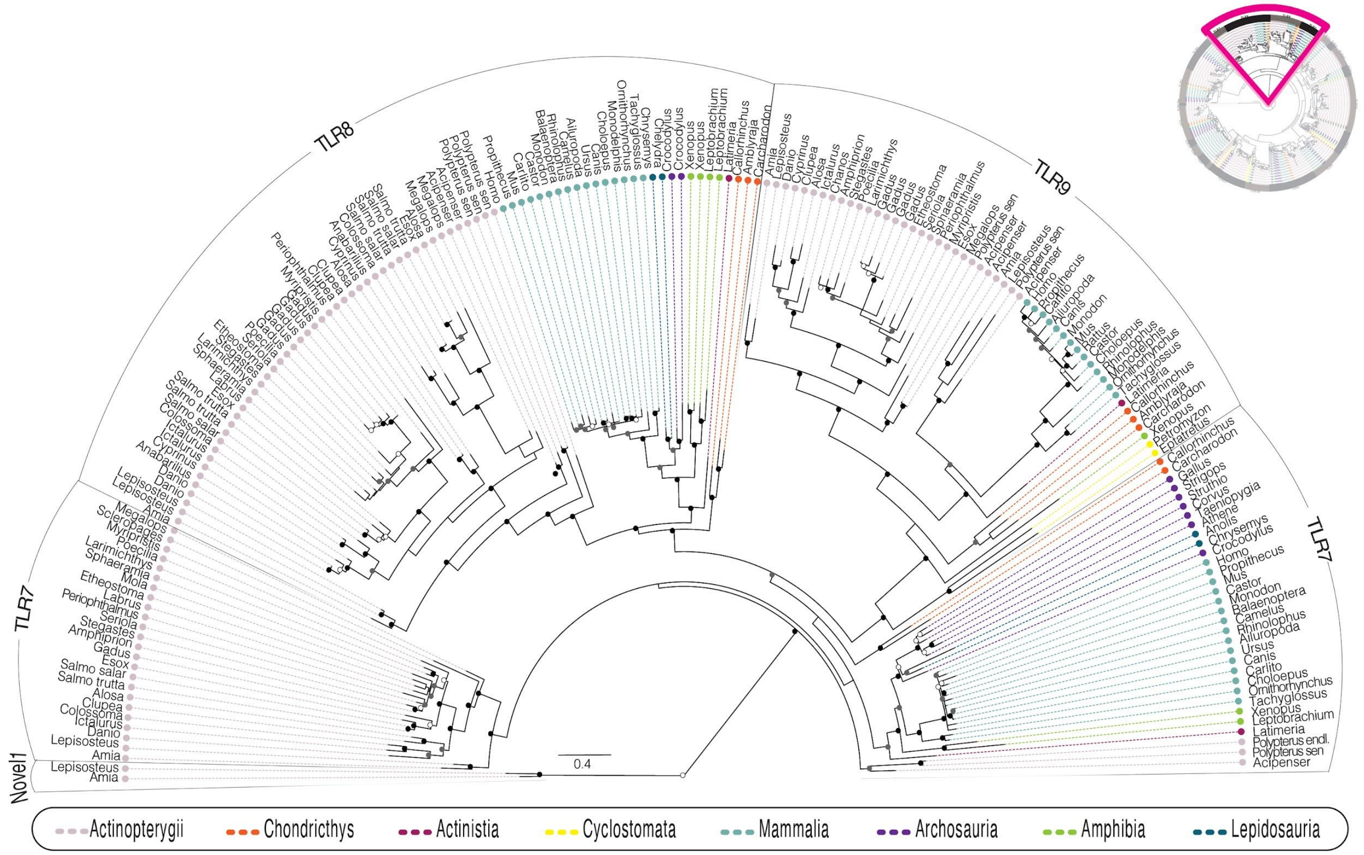

*Supplementary Figure S5. Phylogenetic tree of TLR7 subfamily*

Subtree of **Fig. 1** which shows the relationship of TLR7 subfamilies. Individual sequences are defined by their genus name on the perimeter of the subtree and TLR group designation is indicated by the outermost ring. The position of the subtree within the larger phylogeny (**Fig. 1**) is represented by a pink triangle in the upper right corner. Note the holostean-specific clade of TLRnovel1 sequences. Bootstrap values are represented by black (BSS=100), gray (90 ≤ BSS <100), or white (70 ≤ BSS <90) circles and each class represented by a different color line.

#### ***TLR11 subfamily (11/12/13/19/20/21/22)***

*Supplementary Text 15: TLR11*

TLR11 is a placental mammal-specific TLR, that is present in rodents (*Rattus, Mus, Castor)* as well as tarsier (*Carlito*), sloth (*Choloepus*) and rabbit (*Oryctolagus*) — but not other mammals. Except for a duplication of TLR11 in sloth, TLR11 is a single copy gene in the mammals surveyed here.

*Supplementary Text 16: TLR12*

TLR12 mirrors TLR11 in terms of distribution among taxa, however TLR12 is not seen in tarsier (*Carlito*) *and* is present only as a single copy in sloth (*Choloepus*). A TLR named TLR16 has been annotated in *Xenopus tropicalis* (Roach et al. 2005). — however, in our study we find this TLR to represent the sister lineage to TLR12, TLR11, and TLR19.

*Supplementary Text 17: TLR13*

Our phylogenetic analyses provide strong support for clades of TLR13 that correspond to major vertebrate groups spanning the most recent common ancestor of jawed vertebrates. Although relationships among those groups are not well supported, the strong support for each clade reveals a signal of numerous independent losses. For example, in mammals TLR13 is present in mice but not in rats and present in lemurs but not in tarsiers. TLR13 is also not present in archosaurs and absent from major clades of actinopterygians that include Polypteriformes, Acipenseriformes, Clupeomorpha, and Ostariophysi. In general, TLR13 remains a single copy in all vertebrates with the only notable exceptions within *Megalops* and *Latimeria*. The copies of TLR13 in *Latimeria* are unusual in their sequence divergence and are resolved as belonging in two separate clades with strong support. One of these clades mirrors vertebrate phylogeny. In contrast, the other presents a resolution in which the second copy of TLR13 in *Latimeria* is sister to TLR13 in chondrichthys. This suggests that the second TLR13 in *Latimeria* is not lineage specific, and instead reflects a more ancient gene duplication.

*Supplementary Text 18: TLR19*

TLR19 is a small actinopterygian-specific clade that likely originated prior to the most recent common ancestor of Holostei and Teleostei. TLR19 is maintained in only a few species including *Amia*, *Megalops*, all but one ostariophysan from this study, and both salmon species. All TLR19 sequences except for two found in *Danio* are single copies. TLR19 is absent from acanthomorphs.

*Supplementary Text 19: TLR20*

TLR20 is an actinopterygian-specific clade that likely originated prior to the most recent common ancestor of Holostei and Teleostei. In general, TLR20 is not single copy with few exceptions such as *Alosa* and *Cyprinus*. Although large expansions of TLR20 are found in early diverging euteleosts such as *Salmo* and *Esox*, TLR20 appears to have been lost prior to the most recent common ancestor of acanthomorphs.

*Supplementary Text 20: TLR21*

TLR21 (Supplementary Figure S3) is present in all vertebrates except for mammals. Our phylogenetic analyses provide strong support for several distinct clades of TLR21 lineages. Most of these lineages reflect expectations of general vertebrate phylogeny, such as a clade of Actinopterygian TLR21 and a clade of avian TLR21. However, we find evidence for a deeply divergent clade of TLR21 lineages comprised of *Polypterus*, *Latimeria*, and chondrichthyan lineages. Our results suggest this clade to harbor the vestiges of an early duplication of TLR21 in jawed vertebrates that was independently lost in early sarcopterygian and actinopterygian evolution. During early actinopterygian evolution, our results also suggest a loss of the other TLR21 copy in both Holostei and Acipenseriformes. TLR21 is widespread in teleosts with independent losses in lineages such as *Salmo*.

*Supplementary Text 21: TLR22*

TLR22 arose in the most recent common ancestor of jawed vertebrates. It is present as a single copy gene in some chondrichthyans, amphibians, lepidosaurs, testudines, and crocodilians. However, in actinopterygians TLR22 is often subject to large lineage-specific gene duplications. For example, Colossoma has seven copies of TLR22, and the mudskipper *Periophthalmus* has ten. Our phylogeny suggests that throughout actinopteryigian evolution, some TLR22 paralogs have been maintained and subsequently acted as a genomic substrate for these more recent lineage specific duplications.

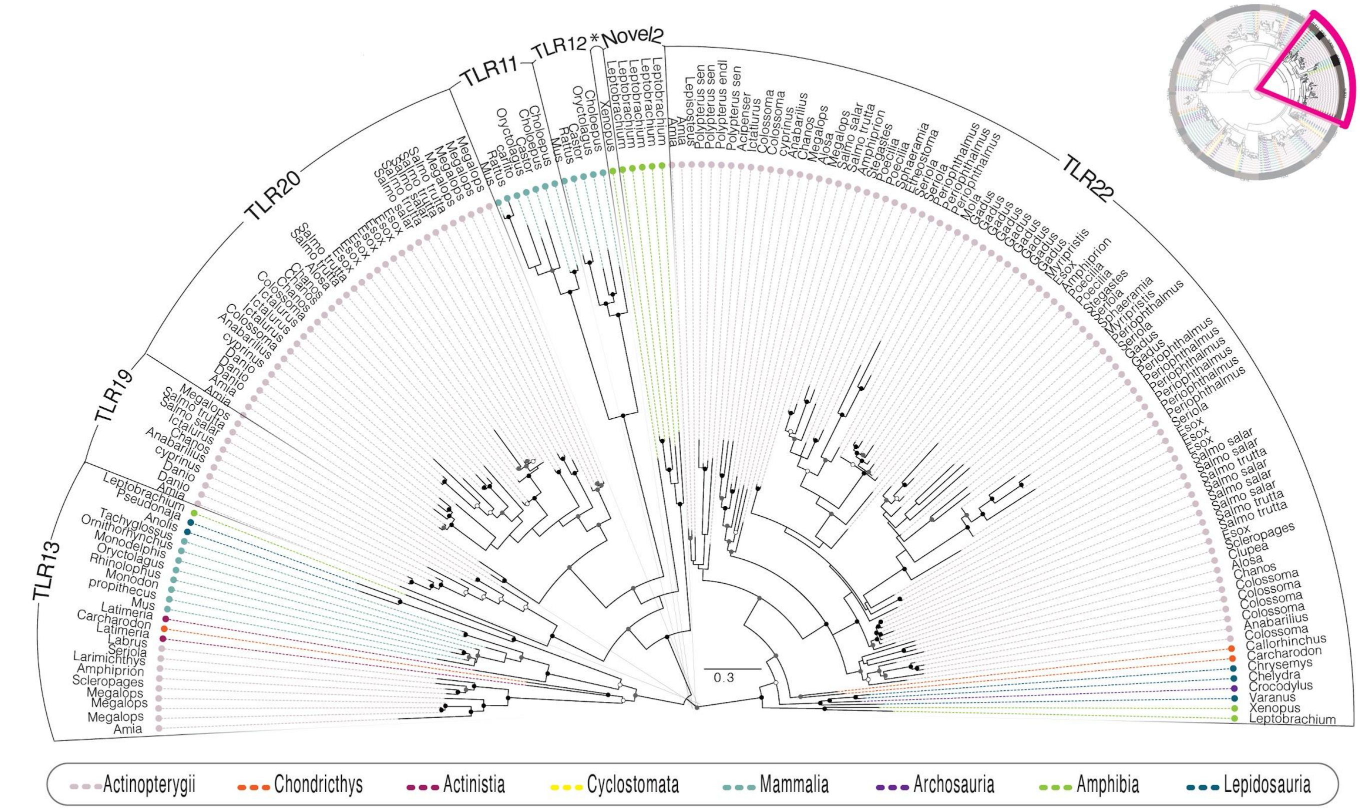

*Supplementary Figure S6. Phylogenetic tree of TLR11 (partial) subfamily*

Subtree of **Fig. 1** which shows the relationship of the TLR11 subfamily (partial). Individual sequences are defined by their genus name on the perimeter of the subtree and TLR group designation is indicated by the outermost ring.The position of the subtree within the larger phylogeny (**Fig. 1**) is represented by a pink triangle in the upper right corner. Note the Leishan spiny toad (*Leptobrachium)*-specific clade of TLRnovel2 sequences. Bootstrap values are represented by black (BSS=100), gray (90 ≤ BSS <100), or white (70 ≤ BSS <90) circles and each class represented by a different color line. The asterisk (*) denotes TLR12 from *Xenopus* that does not fall within any specific clade.

***References***

Berczi I, Bertók L, Bereznai T (1966) Comparative studies on the toxicity of *Escherichia coli* lipopolysaccharide endotoxin in various animal species. *Can J Microbiol* 12(5):1070-1

Bolger AM, Lohse M, Usadel B (2014) Trimmomatic: a flexible trimmer for Illumina sequence data. *Bioinformatics* 30:2114–2120

Carlson KB, Wcisel DJ, Ackerman HD, et al (2022). Transcriptome annotation reveals minimal immunogenetic diversity among Wyoming toads, *Anaxyrus baxteri*. *Conserv Genet* 23, 669–681.

Grabherr MG, Haas BJ, Yassour M, et al (2011) Full-length transcriptome assembly from RNA-Seq data without a reference genome. *Nat Biotechnol* 29:644–652

Jiang W, Lederman MM, Harding CV, Rodriguez B, Mohner RJ Sieg SF (2007) TLR9 stimulation drives naive B cells to proliferate and to attain enhanced antigen presenting function. *Eur. J. Immun.*, *37*: 2205-2213.

Lu YC, Yeh WC Ohashi PS, (2008) LPS/TLR4 signal transduction pathway. *Cytokine*, *42*(2): 145-151.

Manni M, Berkeley MR, Seppey M, et al (2021) BUSCO Update: Novel and Streamlined Workflows along with Broader and Deeper Phylogenetic Coverage for Scoring of Eukaryotic, Prokaryotic, and Viral Genomes. *Mol. Biol. Evol*. 38:4647–4654.

Roach JC, Glusman G, Rowen L, et al (2005) The evolution of vertebrate Toll-like receptors. *Proc Natl Acad Sci U S A* 102:9577–9582
